# Continuum theory of active phase separation in cellular aggregates

**DOI:** 10.1101/2020.02.26.966622

**Authors:** Hui-Shun Kuan, Wolfram Pönisch, Frank Jülicher, Vasily Zaburdaev

## Abstract

Dense cellular aggregates are common in many biological settings, ranging from bacterial biofilms to organoids, cell spheroids and tumors. Motivated by *Neisseria gonorrhoeae* biofilms as a model system, we present a hydrodynamic theory to study dense, active, viscoelastic cellular aggregates. The dynamics of these aggregates, driven by forces generated by individual cells, is intrinsically out-of-equilibrium. Starting from the force balance at the level of individual cells, we arrive at the dynamic equations for the macroscopic cell number density via a systematic coarse-graining procedure taking into account a nematic tensor of intracellular force dipoles. We describe the basic process of aggregate formation as an active phase separation phenomenon. Our theory furthermore captures how two cellular aggregates coalesce. Merging of aggregates is a complex process exhibiting several time scales and heterogeneous cell behaviors as observed in experiments. In our theory, it emerges as a coalescence of active viscoelastic droplets where the key timescales are linked to the turnover of the active force generation. Our theory provides a general framework to study the rheology and dynamics of dense cellular aggregates out of thermal equilibrium.

## Introduction

Complex biological aggregates, such as bacterial colonies, organoids, cell spheroids and tumors, are formed by thousands to millions of individual interacting cells. Studies of dense cellular aggregates provide a bridge between tissues and organs to the level of single cells with implications in various biological contexts. Examples include wound healing [1, 2], tissue spreading [3–6], tumor growth and treatment [7, 8], and formation of biofilms or organoids [8–12]. As the interactions between cells are associated with chemical energy being typically transformed to mechanical work, cellular aggregates are intrinsically out-of-equilibrium (active) systems [13] and behave differently compared to normal viscous liquid droplets.

A prototypical example of active multicellular aggregates are the microcolonies of *Neisseria gonorrhoeae* (*N. gonorrhoeae*) bacteria. *N. gonorrhoeae* microcolonies forming on epithelial tissue of humans are the infectious units of the second most common sexually transmitted diseases gonorrhea. *N. gonorrhoeae* as well as numerous other bacteria species use thin and long retractable filaments called type IV pili to interact with the environment and with each other [14, 15] (see fig. 1 (a)-(c)). Cycles of pili growth, attachment, retraction and detachment drive cell motility on surfaces and the process of colony formation. Forces of actively contracting and constantly remodeling pili networks create a densely packed colony (see fig. 1 (b)) and together with excluded volume interactions form an active viscoelastic material.

**FIG. 1.**
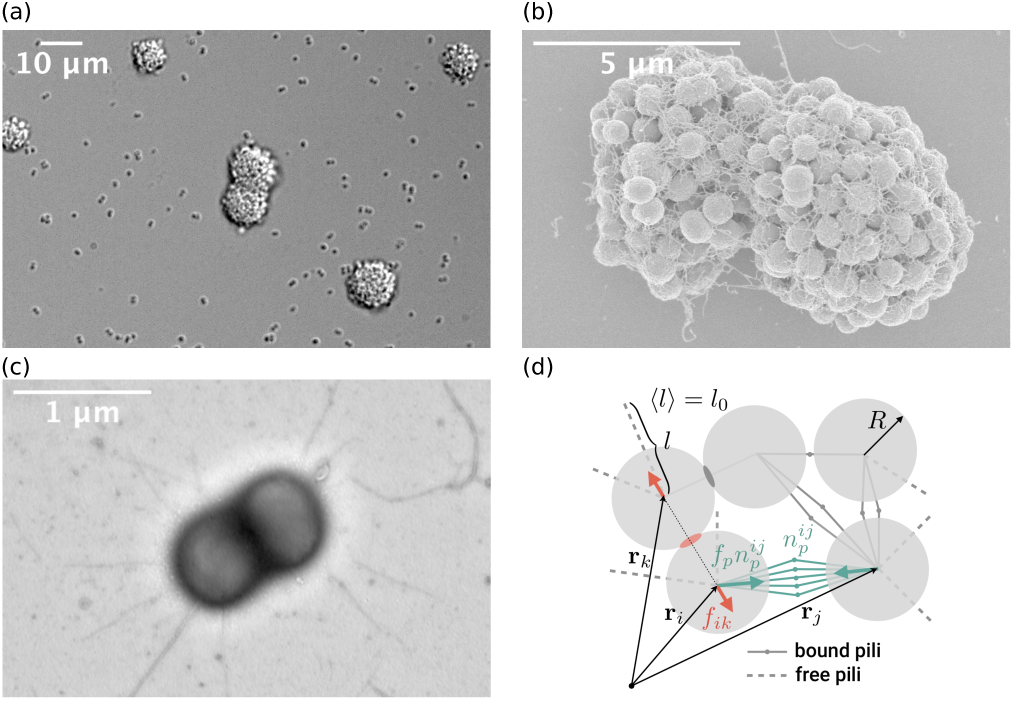
Bacterial colonies and interactions. (a) Optical microscopy images of *N. gonorrhoeae* aggregates formed on a solid substrate. (b) Electron microscopy images of two merging *N. gonorrhoeae* aggregates and pili network. (c) Transmission electron microscopy image of a single *N. gonorrhoeae* cell and its pili. The images (a), (b), and (c) courtesy of N. Biais. (d) Illustration of the microscopic forces (the substrate friction is not shown) acting on the cells. Cells are represented as disks of radius *R*. Solid lines with a dot denote pairs of bound pili. Dashed lines are showing free pili. Pili length *l* is exponentially distributed with the mean length *l*_0_. Cells *i* and *j* have 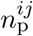 bound pili pairs between them. Each bound pili pair generates a force dipole of strength *f*_p_. Thus, 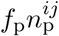 is the magnitude of the force acting from cell *j* on cell *i* due to the pili, while the green arrow shows its direction. *f*_*ik*_ and red arrows show the magnitude and the direction of the steric repulsive force, respectively.

Multi-scale computer simulations have played an important role in the study of the physics of the formation of *N. gonorrhoeae* colonies [16–18]. Continuum approaches such as theories of active gels [19–26] so far do not capture important features of dense cellular aggregates. Here we put forward the continuum theory of dense multicellular systems, where contractile, non-potential, intercellular forces are balanced by excluded volume interactions. We show how the turnover of active forces determines the viscoelastic behavior of aggregates. The developed theory rationalizes the formation of *N. gonorrhoeae* colonies as an active phase separation and highlights the role of viscoelasticity during the complex process of merging between two aggregates. We are thus enabled to theoretically fully understand the rheological behavior of multicellular aggregates.

## Results

### A. Pili-mediated forces

Pili-mediated intercellular interactions of the bacteria are well studied and may serve as an important example to construct a microscopic model for active cellular force generation. *N. gonorrhoeae* bacteria have multiple (10-20) pili isotropically distributed over the cell body (see fig. 1 (c)) [27–29]. Pili grow by a process of polymerization with a speed of ∼ 2 µm*/*s [14, 28, 30, 31]. Pili length is well approximated by an exponential distribution with a mean of *l*_0_ ∼ 1-2 µm (diameter of the cell is ∼ 1 µm) [27], with pili as long as several tens of microns were reported. Pili may attach to a substrate or to pili of other *N. gonorrhoeae* bacteria (but not to their cell membrane) [16]. An attached pilus retracts via depolymerization driven by a specific motor protein complex [32, 33]. The retraction velocity is force dependent, starting from ≈ 2 µm*/*s for an unloaded motor and stalling at the pulling force *f*_p_ ≈ 180 pN [31, 34, 35]. If even a greater stretching force is applied to a pilus, it might extend via a conformational change without detaching [36]. An attached pilus stochastically detaches in a load dependent manner [35]. These are the key pili characteristics which are, in some instances with simplifications, introduced in the microscopic model below.

### B. Cell-based model of aggregate dynamics

In a simplified two-dimensional setting, cells are represented as hard disks with the radius *R* (see fig. 1 (d)). Pili are formed stochastically, with isotropic orientation and with an exponential length distribution with average *l*_0_ [37]. Two individual pili of two neighboring cells may bind to each other. Retraction of the pili will generate no net force acting on the two cells but it will give rise to the attractive force dipole of strength *f*_p_ pulling the cells together (see fig. 1 (d)).

In the overdamped limit, for an individual cell, the pili-pili mediated forces are balanced by excluded volume interactions and substrate friction (see fig. 1 (d)):

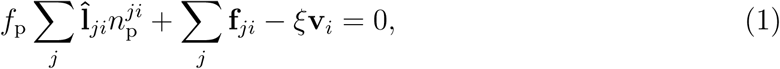

where **v**_*i*_ is the velocity of cell *i* on the substrate and *ξ* is a friction coefficient. The steric repulsion force exerted by cell *j* on cell *i* is denoted **f**_*ji*_. The unit vector 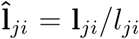 shows the direction of the pili-pili mediated force exerted by cell *j* on cell *i*, where **l**_*ji*_ = **r**_*j*_ − **r**_*i*_ is the vector pointing from cell *i* at the position **r**_*i*_ to cell *j* at the position **r**_*j*_, and *l*_*ji*_ is the corresponding distance. The pili-pili mediated force between cells is proportional to the number of bound pili pairs between cells *i* and *j*, denoted 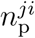 (see fig. 1 (d)). Note that 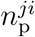 is changing with time. We consider formation of bound pili pairs to occur with a binding constant *k*_on_ and the detachment with a constant unbinding rate *k*_off_. Because the steady-state pili length distribution is exponential, the number of bound pili pairs between cells *i* and *j* obeys (for details see the supplementary information [SI]):

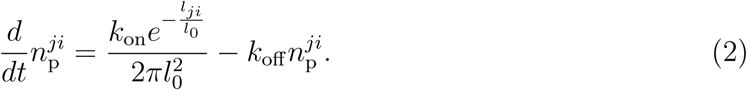

The total number of bound pili of an individual cell *i* can be obtained as the sum over all bound pili pairs with all of its neighboring cells *j* and thus in general depends on the cell number density and binding/unbinding constants. However, biologically, the total number of pili per cell could have an upper limit [16, 27, 38, 39]. This limitation can be incorporated in Eq. (2) (see the supplementary information [SI]) but which we forego for the sake of simplicity.

### C. Continuum limit

The next conceptual step is to use the microscopic force balance equation together with pili turnover dynamics to obtain dynamic equations for coarse-grained cell number density and cell velocity field. The coarse-graining procedure is described in the supplementary information [SI]. There are two key features of the considered system, which distinguish it from previous models of active particle systems [40–43]. First, the network of bound pili leads to an elastic-like response of the colony when it is deformed at time scales smaller than the pili detachment time: attached pili stretch at constant force. At larger time scales, pili can detach and reattach in a new configuration which leads to stress relaxation and thus a more fluid-like material response. Second, the active pili-pili mediated forces between two cells act as attractive force dipoles. These active attractive interactions are balanced by steric repulsion forces when cells get in direct contact and are pushed against each other forming a dense cellular aggregate.

These two features as well as details of microscopic interactions of cells are fully captured by the continuum equations (for derivation see the supplementary information [SI]) for the cell number density *n* and force balance:

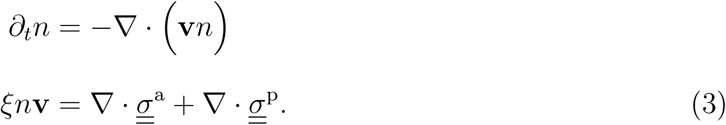

Here **v** is the cell velocity field, and 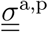 are the active and passive stress tensors, respectively.

The expression for the passive stress incorporates effects from short range repulsive interactions. In the continuum, we use the expression

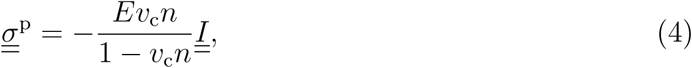

where *v*_c_ = *πR*^2^ is the area of a single cell, *E* is the bulk elastic modulus, and 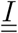 is the identity matrix. This, in fact, is the van der Waals gas law for pressure without attractive interaction, and is one of the simplest forms to describe the excluded volume effects.

The active stress is related to the dynamics of bound pili. Due to the intrinsic nature of pili-pili interaction as force dipoles [23], the first non-trivial term of the active stress has nematic symmetry [23, 44, 45] and can be written as

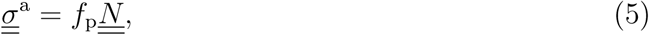

where 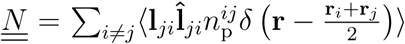 is a nematic tensor which characterizes the amount of the bound pili pairs and their axis of anisotropy, *δ* (…) is the Dirac delta function, and ⟨*…* ⟩ denotes the ensemble average (see the the supplementary information [SI]).

### D. Phase separation

The continuum theory can give rise to a phase separation regime where attractive pili-pili forces drive the formation of dense aggregates from an initially homogneous distribution of cells (see fig. 2 (a) and (b) and supplementary movie). In the case where the dynamics of bound pili pairs in Eq.(2) relaxes faster than the time required for the bacteria to move over the distance of its own radius, we can use the adiabatic approximation for the active stress. In this case, we can find a relation that determines the critical density *n*_*c*_ above which the initially homogeneous state of the system becomes unstable with respect to small density fluctuations. This critical density *n*_*c*_ obeys (fig. 2 (c) and the supplementary information [SI]):

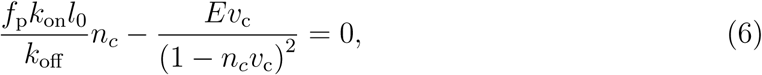

which follows from linear stability analysis. Beyond this critical density, cells form dense aggregates.

**FIG. 2.**
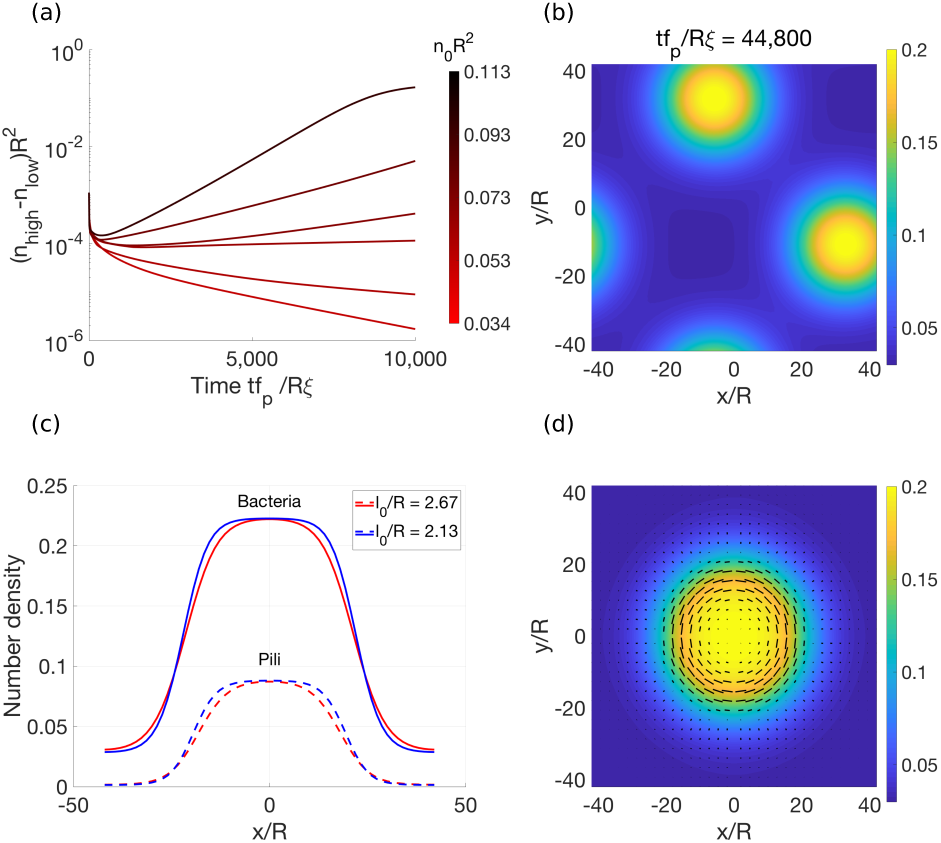
Aggregate formation results from a phase separation phenomenon in an active system. (a) Density difference of high and low cell number density phases as a function of time. The critical initial density of the onset of the active phase separation is at *n*_*c*_*R*^2^ ≃ 0.049 for the ratio *k*_on_*/k*_off_ *R*^2^ = 1.78, and the initial density fluctuation level of 0.001. (b) The snapshots of the late time of the phase separation. The characteristic pili length is *l*_0_*/R* = 2.67, the initial density is *n*_0_*R*^2^ = 0.068, and *f*_p_*/πRE* = 6. (c) The cell number density (solid lines) and total number of bound pili density (dashed lines) of stable droplets (aggregates) for two different characteristic pili lengths. The bulk elastic moduli are taken as *f*_p_*/πRE* = 6 (red) and *f*_p_*/πRE* = 7.5 (blue) to provide the same degree of phase separation. The characteristic pili length controls the width of the aggregate’s boundary. The initial density for this graph is *n*_0_*R*^2^ = 0.072. (d) The nematic tensor 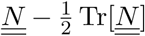 inside of a stablized aggregate. The initial density is *n*_0_*R*^2^ = 0.072, the characteristic pili length is *l*_0_*/R* = 2.67. Black lines indicate the nematic ordering direction, and their length shows the nematic order parameter.

### E. Viscoelastic dynamics

We next discuss the dynamics of colony formation, which is strongly dependent on pili dynamics. The material properties of colonies mediated by pili forces are controlled by the nematic tensor 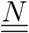. In order to describe the dynamics of colonies, we determine the time evolution of the active stress in our coarse-grained theory and show that it corresponds to a viscoelastic behavior (see the supplementary information [SI]). Calculating the time derivative of the active stress tensor naturally leads to the appearance of higher rank tensors. We estimate these higher rank terms using a standard closure approximation, and introducing the density of the number of bound pili *N* (see the supplementary information [SI]). The active stress then obeys

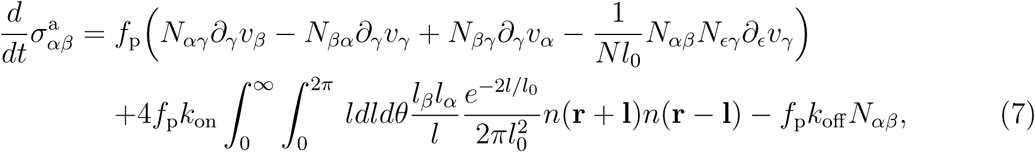

where the tensorial notation with Greek letters indicate the Cartesian coordinates and the repeated Greek indices indicate Einstein’s summation convention. The active stress relaxes at a rate *k*_off_, which implies viscoelastic material properties. Equation (7) describes the stress–strain relation due to cell movement (first parenthesis), the active stress from the newly formed pili pairs (second term), and the relaxation of the active stress from the detachment of bound pili (last term). Combined with the definition of the active stress (Eq. (5)), the relaxation dynamics can be written as the Maxwell model with a source term:

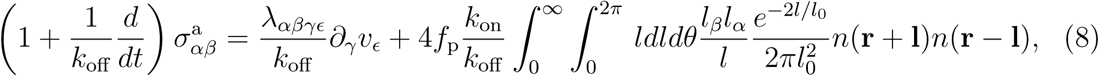

Where 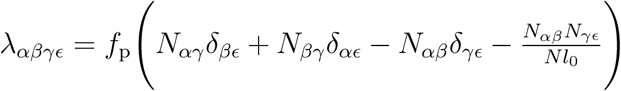 is an elastic tensor. At short times 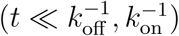, the active stress behaves elastically. At larger times 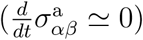, the network of bound pili pairs is remodeled due to the binding/unbinding processes and the active stress behaves as in a viscous fluid with the viscous tensor 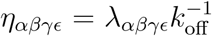 (calculated from the ratio between the elastic strain and relaxing stress).

In our model, pili-induced elastic moduli and viscosities explicitly depend on the cell number density. In the boundary region (the surface of the colony), the cell number density is lower than in the bulk and thus we expect the elastic moduli and the viscosities to be smaller at the surface. Thus our theory naturally accounts for the previously experimentally observed effect of heterogeneous cell motility in *N. gonorrhoeae* aggregates with higher and lower cell dynamics at the surface layer and in the bulk of the colony, respectively [16]. This heterogeneity in turn effects the merging of colonies, a complex dynamic process and the major mechanism of colony growth in *N. gonorrhoeae*, which we consider next.

### F. Coalescence of colonies

The merging of cellular aggregates often is reminiscent of a coalescence of liquid droplets [16, 46–48]. One way to quantify this process is to follow the time dynamics of the liquid (capillary) bridge — the contact area of the droplets. We can measure the bridge height *h*(*t*) (see fig. 3 (a)) as a function of time to quantify the progression of merging. Overall we see an interesting behavior which is not captured by a single time scale. Because of the viscoelastic nature of the active stress (see Eq.6), at short times 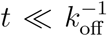, the merging is dominated by the cell movements without detachment of the bound pili, which leads to an elastic response with an effective elastic modulus given above. At large times 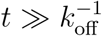, the bound pili network has enough time to remodel and leads to an effective viscous behavior and gives rise to a time scale governed by the ratio of the effective viscosity to the average stresses.

**FIG. 3.**
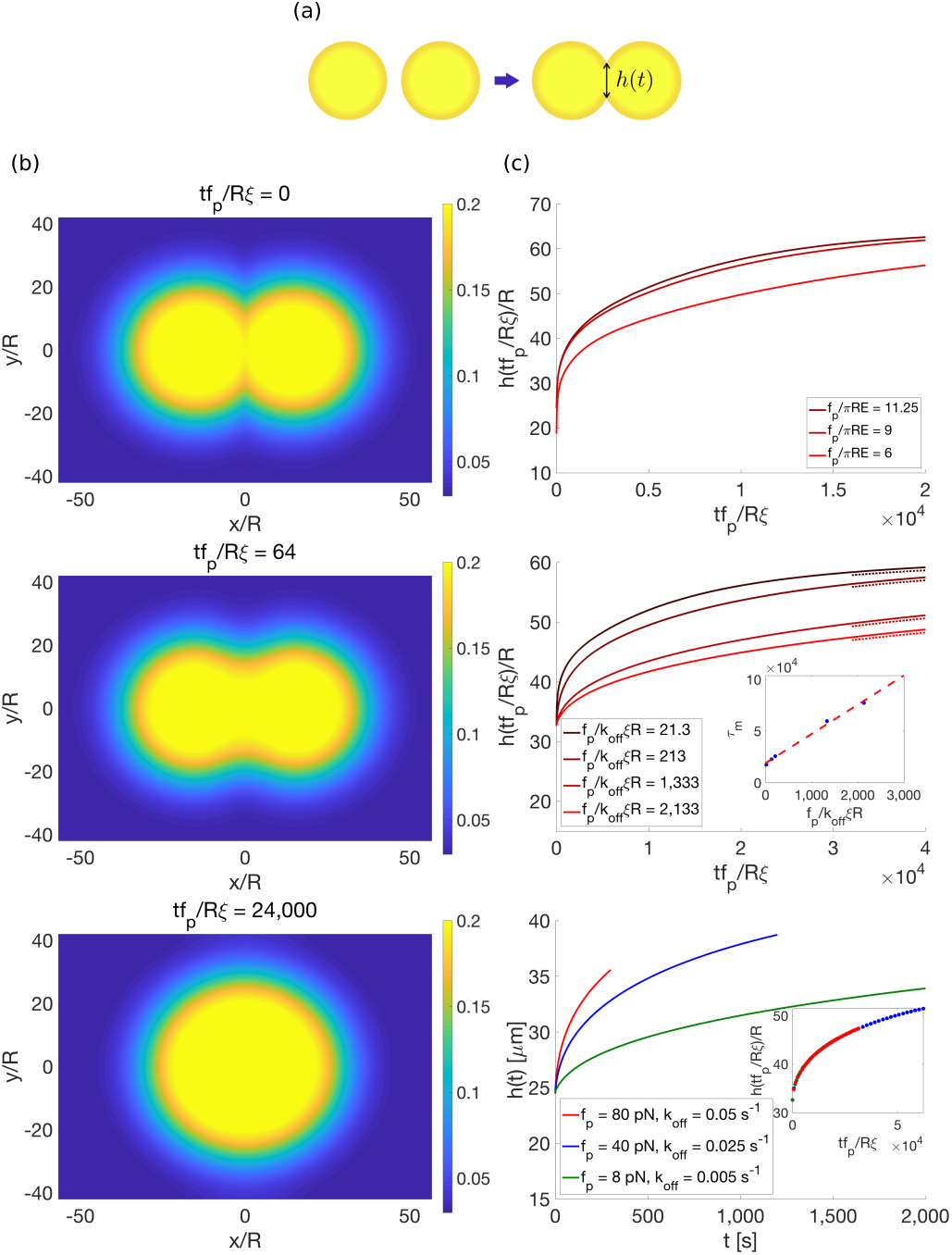
Coallescence of aggregates. (a) Definition of the bridge height *h*(*t*). The boundaries of the merged aggregate are defined at the mid point between the low and high density levels of the initial aggregates. (b) The snapshots of two aggregates merging at different times. The binding rate constant is *k*_on_*ξ/f*_p_*R* = 0.083, unbinding rate constant is *k*_off_ *ξR/f*_p_ = 0.047, and *f*_p_*/πRE* = 6. (see supplementary movie) (c) The bridge height *h*(*tf*_p_*/Rξ*)*/R* as the function of the rescaled time *tf*_p_*/Rξ*. Top: the bridge height for different values of *f*_p_, while *k*_on_ and *k*_off_ are kept constant. The gradient of colors indicates the values of *f*_p_, with darker color corresponding to higher force. Middle: the bridge height for varying *k*_off_, where *f*_p_*/πRE* = 6, however, also the ratio *k*_on_*/R*^2^*k*_off_ = 1.78 is kept constant to guarantee the same degree of phase separation. The dashed lines are the late time fitting (for details see the supplementary information [SI]) with a constant shift in height for the sake of clarity. The gradient of colors indicates the values of *k*_off_, with the darker color corresponding to higher value of *k*_off_. The inset shows the asymptotic merging time *τ*_*m*_ (corresponding to the dashed lines in the main plot) which linearly depends on 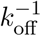 due to the effective viscosity. Bottom: the bridge height for different numerical values of *k*_off_ and *f*_p_. All these curves have the fixed values of 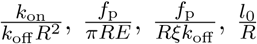 and *ξ =* 1*pNs/*μm, while the inset shows the master curve with the rescaled time *tf*_p_*/Rξ*. The red curve is close to the parameter regime considered in the previous study [16].

With the help of the model we can now test how the merging dynamics depends on the relevant parameters. By varying the strength of force dipole *f*_p_ and the pili detachment rate *k*_off_, aggregates tend to merge faster for increasing *f*_p_ and decreasing *k*_off_ (see fig.3(c)). Because of the viscoelastic nature of the active stress (see Eq.6), the system behaves like a viscoelastic active liquid and an additional time scale from ratio of the effective viscosity and the stress tensor become relevant during the late stage of the merging [49–52]. The merging time scale at the late stage can be shown to linearly depend on 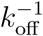 (see the inset in the middle panel of fig.3 (c)) due to the scaling of the effective viscosity with 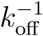. By dimensional analysis, we find that the system depends on four non-dimensional parameters: 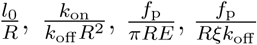 and on the rescaled time 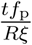 when the initial cell number density is fixed. The first parameter is the ratio of the mean pili length and the cell radius, the second parameter is related to the number of bound pili, and the third parameter is the ratio between the active force and excluded volume force. These three non-dimensional parameters determine the degree of the active phase separation (see also Eq.6). The fourth non-dimensional parameter is the ratio between the time of moving a cell half of its size under the pili force and the pili detachment time, which is the only non-dimensional parameter which we can vary if the degree of the phase separation is fixed. The fig.3 (c) (bottom) confirms these scalings. The plots showing bridge height as a function of time describe merging process for different parameters collapse on a single master curve when we apply the appropriate scaling of length and time.

Our work provides a systematic derivation of a continuum theory to study active phase separation and the active rheology of dense cellular aggregates. The contribution of active intercellular pili-mediated forces is incorporated via binding and detachment kinetics. Force generation similar to pili-induced active stresses is common in many biological systems, not only interacting cells but for example also in active gels [23, 53]. The remodeling of the active force gives rise to time dependent or permanent (if the active force persists) rheological responses. In our case, relaxation of active stress proceeds through the detachment of bound pili. In other situations it may occur through the relaxation of nematic order [23, 24, 44]. The coupling between the active force generation and cell motion gives rise to space and time dependent elastic moduli and viscosities, which makes our model suitable to treat imhomogeneous systems, a prerequisite to eventually understand morphogenesis of complex tissues.

#### Numerical methods and parameter choices

We use XMDS2 package [54] with eighth- and nineth-order adaptive Runge-Kutta method to numerically solve the differential equations (see the supplementary information [SI] equations (S34)). The simulations for active phase separation (fig. 2(a)) are performed with the parameters *f*_p_*/πER* = 6, *k*_on_*/k*_off_ *R*^2^ = 1.78, *l*_0_*/R* = 2.67, and with the steady-state nematic tensor at any time step during the evolution (the adiabatic approximation). We have tested several cases to prove the non-steady-state nematic tensor assumption won’t change the results at the steady state. The well-developed aggregates are prepared in the two-dimensional periodic box with 64 *×* 64 bins with the initial density *n*_0_*R*^2^ = 0.072. The aggregates are considered well-developed aggregates when the change of the radii of the aggregates are smaller than 1% in the time steps longer than 1 *×* 10^4^ (in the unit of *Rξ/f*_p_) with the assumption of the steady-state nematic tensor at any time step. The initial conditions for the merging in Fig. 3 (c) are prepared by horizontally putting two boxes of the well-developed aggregates together. In both boxes, we cut 21 bins (it is 17 bins in the upper panel of Fig. 3(c)) on the sides with which aggregates are facing each other horizontally, so that in the resulting simulations on box of 64 *×* 86 bins (64 *×* 94 in the upper panel of Fig. 3(c)) two aggregates are touching each other. The merging times in the inset of the middle panel of Fig. 3 (c) are from fitting the last one-forth of the bridge height (see the main text) with the function: *h*(*t*) = *h*_*∞*_ (1 *-a* exp(*-t/τ*_*m*_)), where *τ*_*m*_ is defined as the merging time, and *h*_*∞*_ ∼ 60.58 (in the unit of *R*) is the final diameter of the merging.

## Acknowledgments

We thank Christoph A. Weber, Taylor Harmon, Jakob Löber, Suropriya Saha, Lennart Hilbert, and Rabea Seyboldt for useful discussions on the concepts of phase separation and assistance with numerical modeling. We thanks Nicolas Biais for detailed discussion and providing images. This work was supported by Volkswagen foundation “Life?” initiative (Hui-Shun Kuan and Vasily Zaburdaev). Wolfram Pönisch thanks the Leverhulme Trust and the Herchel Smith Postdoctoral Fellowship Fund.

## Author contributions

This was a collaboration of all authors. All authors designed the work and wrote the paper. Hui-Shun Kuan performed analytical calculations and numerical simulations.

