## Supplementary information for "Continuum theory of active phase separation in cellular aggregates"

### I. THE CONTINUUM MODEL

To build the continuum model of the pili-induced formation of bacterial aggregates in two dimensions, we start from the microscopic description and derive the continuum model via the coarse-graining.

The bacterium $i$ at position $\mathbf{r}_i$ has velocity $\mathbf{v}_i(t)$:

$$\frac{d}{dt}\mathbf{r}_i(t) \equiv \mathbf{v}_i(t). \tag{S1}$$

In the overdamped limit, for each bacterium $i$, the force balance equation including the pili-induced force, the excluded volume force, and the friction reads:

$$f_{\mathrm{p}} \sum_{j \neq i} \hat{\mathbf{l}}_{ji} n_{\mathrm{p}}^{ji} + \sum_{j \neq i} \mathbf{f}_{ji} - \xi \mathbf{v}_i = 0. \tag{S2}$$

If two bacteria $i$ and $j$ have $n_{\mathrm{p}}^{ij}$ number of bound pili pairs between them, there is a force $f_{\mathrm{p}}\hat{\mathbf{l}}_{ji} n_{\mathrm{p}}^{ji}$ acting on bacterium $i$ in the direction $\hat{\mathbf{l}}_{ji} \equiv \frac{\mathbf{l}_{ji}}{|\mathbf{l}_{ji}|} = \frac{\mathbf{r}_j - \mathbf{r}_i}{|\mathbf{r}_j - \mathbf{r}_i|}$, where $f_{\mathrm{p}}$ is the force generated by a single pulling pair of bound pili. For the force balance we sum over all bacteria $j$ interacting with the bacterium $i$. $\mathbf{f}_{ij}$ is the excluded volume force between two different bacteria $i$ and $j$. $\xi \mathbf{v}_i$ is the friction force with an effective friction coefficient $\xi$. Friction in our model is a combined effect of solvent, surface and surrounding bacteria. Its main phenomenological purpose is to balance pili pulling force and produce constant speed motion, which in the bacterium is advised by the constant speed of pili retraction [1].

The number of bound pili pairs $n_{\mathrm{p}}^{ji}$ between bacteria $i$ and $j$ is controlled by the binding and

*

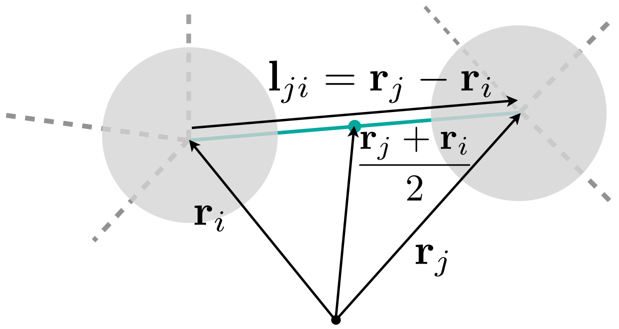


FIG. S1. The illustration of the bacteria and bound pili. The bacteria are labelled as gray circles, the green lines are the bound pili, and the dashed gray lines are pili. The bacteria are in the position $\mathbf{r}_j$ and $\mathbf{r}_i$, and we denote the bound pili position (green circle) as the midpoint between the two bacteria :$\frac{\mathbf{r}_j+\mathbf{r}_i}{2}$. The vector $\frac{\mathbf{l}_{ji}}{2} = \frac{\mathbf{r}_j-\mathbf{r}_i}{2}$ denotes the bound pili length and the orientation.

unbinding processes:

$$\frac{d}{dt} n_{\mathrm{p}}^{ji} = \frac{k_{\mathrm{on}} e^{-\frac{l_{ji}}{l_0}}}{2\pi l_0^2} - k_{\mathrm{off}} n_{\mathrm{p}}^{ji}, \tag{S3}$$

where $k_{\mathrm{on}}$ and $k_{\mathrm{off}}$ are the binding and unbinding rate constants. The factor $\frac{k_{\mathrm{on}} e^{-l_{ji}/l_0}}{2\pi l_0^2}$ comes from the assumption of an exponential distribution of the pili length and integration over all possible binding points along the line connecting two bacteria. Since $l_{ji} =\mid \mathbf{l}_{ji} \mid$ is the distance between the two bacteria $i$ and $j$, $n_{\mathrm{p}}^{ji}$ has the symmetry $n_{\mathrm{p}}^{ji} = n_{\mathrm{p}}^{ij}$.

#### A. The coarse-grained representation

The coarse-grained representation can be derived by summation over all bacteria at a given position $\mathbf{r}$. The the number density, $n(\mathbf{r}, t)$, can be written as:

$$n(\mathbf{r}) = \sum_i \left\langle \delta\Big(\mathbf{r} - \mathbf{r}_i(t)\Big) \right\rangle, \tag{S4}$$

where $\langle \cdots \rangle$ denotes the ensemble average. Then, for any given physical microscopic quantities $A_i$, and $A_i B_i$, the corresponding coarse-grained values $A(\mathbf{r})$, and $A(\mathbf{r}, t)B(\mathbf{r}, t)$, are defined as:

$$\begin{aligned} \sum_i \left\langle A_i \delta\Big(\mathbf{r} - \mathbf{r}_i(t)\Big) \right\rangle &\equiv n(\mathbf{r}, t) A(\mathbf{r}, t) \\ \sum_i \left\langle A_i B_i \delta\Big(\mathbf{r} - \mathbf{r}_i(t)\Big) \right\rangle &\simeq \frac{\sum_i \left\langle A_i \delta\Big(\mathbf{r} - \mathbf{r}_i(t)\Big) \right\rangle \sum_i \left\langle B_i \delta\Big(\mathbf{r} - \mathbf{r}_i(t)\Big) \right\rangle}{\sum_i \left\langle \delta\Big(\mathbf{r} - \mathbf{r}_i(t)\Big) \right\rangle} \\ &= n(\mathbf{r}, t) A(\mathbf{r}, t) B(\mathbf{r}, t). \end{aligned} \tag{S5}$$

The coarse-grained values of the cross physical quantities: $A_{ij}$, $A_i B_j$, and $A_{ij} B_j$ are defined as:

$$\sum_{i \neq j} \left\langle A_{ij} \delta\Big(\mathbf{r} - \mathbf{r}_i(t)\Big) \delta\Big(\mathbf{r}' - \mathbf{r}_j(t)\Big) \right\rangle \simeq n(\mathbf{r}, t) n(\mathbf{r}', t) A(\mathbf{r}, \mathbf{r}', t)$$

$$\sum_{i\neq j}\Big\langle A_iB_j\delta\Big(\mathbf{r}-\mathbf{r}_i(t)\Big)\delta\Big(\mathbf{r}'-\mathbf{r}_j(t)\Big)\Big\rangle \simeq \sum_i\Big\langle A_i\delta\Big(\mathbf{r}-\mathbf{r}_i(t)\Big)\Big\rangle\sum_j\Big\langle B_j\delta\Big(\mathbf{r}'-\mathbf{r}_j(t)\Big)\Big\rangle$$
$$= n(\mathbf{r},t)n(\mathbf{r}',t)A(\mathbf{r},t)B(\mathbf{r}',t)$$
$$\sum_{i\neq j}\Big\langle A_{ij}B_j\delta\Big(\mathbf{r}-\mathbf{r}_i(t)\Big)\delta\Big(\mathbf{r}'-\mathbf{r}_j(t)\Big)\Big\rangle \simeq \frac{\sum_{i\neq j}\Big\langle A_{ij}\delta\Big(\mathbf{r}-\mathbf{r}_i(t)\Big)\delta\Big(\mathbf{r}'-\mathbf{r}_j(t)\Big)\Big\rangle\sum_j\Big\langle B_j\delta\Big(\mathbf{r}'-\mathbf{r}_j(t)\Big)\Big\rangle}{\sum_j\Big\langle\delta\Big(\mathbf{r}'-\mathbf{r}_j(t)\Big)\Big\rangle}$$
$$= n(\mathbf{r},t)n(\mathbf{r}',t)A(\mathbf{r},\mathbf{r}',t)B(\mathbf{r}',t). \tag{S6}$$

It is worth to note that the coarse-grained values $A(\mathbf{r})$, $A(\mathbf{r},\mathbf{r}')$, $\cdots$ can be understood as the average of their microscopic quantities $A_i$, $A_{ij}$, $\cdots$ at the position $\mathbf{r}$, or positions $\mathbf{r}$ and $\mathbf{r}'$, respectively.

Then, according to the above rules, the number density continuity equation can be derived from the microscopic equation (S1):

$$\partial_t n(\mathbf{r},t) = \sum_i -\Big\langle\mathbf{v}_i(t)\cdot\nabla\delta\Big(\mathbf{r}-\mathbf{r}_i(t)\Big)\Big\rangle = -\nabla\cdot\Big(\mathbf{v}(\mathbf{r},t)n(\mathbf{r},t)\Big). \tag{S7}$$

The continuous version of the force balance equation (S2) is then written as:

$$\sum_i\left\{\sum_{j\neq i}\Big\langle f_\mathrm{p}\hat{\mathbf{l}}_{ji}n_\mathrm{p}^{ji}\delta\Big(\mathbf{r}-\mathbf{r}_i(t)\Big)\Big\rangle+\sum_{j\neq i}\Big\langle\mathbf{f}_{ji}\delta\Big(\mathbf{r}-\mathbf{r}_i(t)\Big)\Big\rangle-\Big\langle\xi\mathbf{v}_i\delta\Big(\mathbf{r}-\mathbf{r}_i(t)\Big)\Big\rangle\right\}$$
$$= \mathbf{f}_\mathrm{p}+\mathbf{f}_\mathrm{ex}-\xi n\mathbf{v} = 0. \tag{S8}$$

Here we make a summation over all bacteria $i$ at position $\mathbf{r}$, and the corresponding force densities are $\mathbf{f}_\mathrm{p}$, which is the coarse-grained pili-induced force density, $\mathbf{f}_\mathrm{ex}$, which denotes the excluded volume interaction, and the friction $-\xi n\mathbf{v}$.

Both the excluded volume interaction and the pili-induced force density are related to two interacting bacteria $i$ and $j$. To facilitate calculations we introduce an additional delta function by coarse-graining over the bacteria $j$ at the position $\mathbf{r}'$. The pili-induced and excluded volume interaction force densities then can be written as:

$$\mathbf{f}_\mathrm{p} = \int d^2r'\sum_{i\neq j}\Big\langle f_\mathrm{p}\hat{\mathbf{l}}_{ji}n_\mathrm{p}^{ji}\delta\Big(\mathbf{r}-\mathbf{r}_i(t)\Big)\delta\Big(\mathbf{r}'-\mathbf{r}_j(t)\Big)\Big\rangle$$
$$\mathbf{f}_\mathrm{ex} = \int d^2r'\sum_{i\neq j}\Big\langle\mathbf{f}_{ji}\delta\Big(\mathbf{r}-\mathbf{r}_i(t)\Big)\delta\Big(\mathbf{r}'-\mathbf{r}_j(t)\Big)\Big\rangle, \tag{S9}$$

where integration is over all space. By utilizing the symmetries $\mathbf{f}_{ji}=-\mathbf{f}_{ij}$ and $\hat{\mathbf{l}}_{ji}n_\mathrm{p}^{ji}=-\hat{\mathbf{l}}_{ij}n_\mathrm{p}^{ij}$ and performing the Taylor expansion around the midpoint between two bacteria at $\frac{\mathbf{l}_{ji}}{2}$, we can relate the force densities and the corresponding stress tensors:

$$\mathbf{f}_\mathrm{p} \simeq \int d^2r'\sum_{i\neq j}\left\langle f_\mathrm{p}\hat{\mathbf{l}}_{ji}n_\mathrm{p}^{ji}\delta\Big(\mathbf{r}'-\mathbf{r}_j(t)-\frac{\mathbf{l}_{ji}}{2}\Big)\frac{\mathbf{l}_{ji}}{2}\cdot\nabla_\mathbf{r}\delta\Big(\mathbf{r}-\mathbf{r}_i(t)-\frac{\mathbf{l}_{ji}}{2}\Big)\right\rangle$$
$$\equiv \nabla\cdot\underline{\underline{\sigma}}^\mathrm{a}, \text{ and}$$
$$\mathbf{f}_\mathrm{ex} \simeq \int d^2r'\sum_{i\neq j}\left\langle\mathbf{f}_{ji}\delta\Big(\mathbf{r}'-\mathbf{r}_j(t)-\frac{\mathbf{l}_{ji}}{2}\Big)\frac{\mathbf{l}_{ji}}{2}\cdot\nabla_\mathbf{r}\delta\Big(\mathbf{r}-\mathbf{r}_i(t)-\frac{\mathbf{l}_{ji}}{2}\Big)\right\rangle \tag{S10}$$
$$\equiv \nabla\cdot\underline{\underline{\sigma}}^\mathrm{p}, \text{ and,} \tag{S11}$$

where $\underline{\underline{\sigma}}^{\rm p}$ is the passive stress tensor and $\underline{\underline{\sigma}}^{\rm a}$ denotes the active stress tensor. Below we will make use of the fact that the integration of the product of two delta functions can be rewritten by changing the variable $\mathbf{r}' = \mathbf{r} + 2\mathbf{l}$:

$$\begin{aligned}
&\int d^2r'\delta\Big(\mathbf{r} - \mathbf{r}_i(t) - \frac{\mathbf{l}_{ji}}{2}\Big)\delta\Big(\mathbf{r}' - \mathbf{r}_j(t) - \frac{\mathbf{l}_{ji}}{2}\Big)\\
&= \int 4d^2l\delta\left(\mathbf{r} - \frac{\mathbf{r}_i(t)+\mathbf{r}_j(t)}{2}\right)\delta\left(2\mathbf{l} - (\mathbf{r}_j(t) - \mathbf{r}_i(t))\right), \text{or} &\text{(S12)}\\
&= \delta\left(\mathbf{r} - \frac{\mathbf{r}_i(t)+\mathbf{r}_j(t)}{2}\right) &\text{(S13)}
\end{aligned}$$

where $2\mathbf{l}$ is the difference vector between the two bacteria $i$ and $j$.

#### B. The active stress tensor

The active stress tensor (eq. (S11)) rooted in active dipole forces of bound pili maybe viewed as the nematic tensor $N_{\alpha\beta}$ multiplied by the amplitude of the pulling force $f_{\rm p}$:

$$\begin{aligned}
\sigma^{\rm a}_{\alpha\beta} &= f_{\rm p}N_{\alpha\beta}, \text{and} &\text{(S14)}\\
N_{\alpha\beta} &= \int 4d^2l\sum_{i\neq j}\frac{l_\alpha l_\beta}{l}\left\langle n_{\rm p}^{ij}\delta\left(\mathbf{r} - \frac{\mathbf{r}_i(t)+\mathbf{r}_j(t)}{2}\right)\delta\left(2\mathbf{l} - (\mathbf{r}_j(t) - \mathbf{r}_i(t))\right)\right\rangle,
\end{aligned}$$

where the tensorial notation with Greek letters which indicate the Cartesian coordinates has been used, and the repeated Greek indices indicate Einstein's summation convention. In this article, the tensorial notation for any vector is $\mathbf{A}_i = (A^i_x, A^i_y)$.

The active stress tenor (or the nematic tensor) changes in time due to the pili-binding/unbinding process and relative motion of the bacteria. Here we refer to the experimentally known fact that when pili are pulled on with a force exceeding the stalling force of the motor responsible for pili retraction, they can go through the conformational change allowing them to extend without rapturing or detaching [2]. This effect would correspond to a constant force, which does not depend on the relative distance between bacteria if the pili are bound. The time evolution of the active stress tensor thus can be written as:

$$\begin{aligned}
\partial_t\sigma^{\rm a}_{\alpha\beta} \simeq f_{\rm p}\int 4d^2l\sum_{i\neq j}\Bigg\langle\frac{l^{ji}_\alpha}{l_{ji}}n_{\rm p}^{ij}\Bigg[&\left(\frac{v^j_\beta - v^i_\beta}{2}\delta\left(\mathbf{r} - \frac{\mathbf{r}_i(t)+\mathbf{r}_j(t)}{2}\right)\delta\left(2\mathbf{l} - (\mathbf{r}_j(t) - \mathbf{r}_i(t))\right)\right)\\
&-\partial_\gamma\left(\frac{l^{ji}_\beta}{2}(\frac{v^j_\gamma + v^i_\gamma}{2})\delta\left(\mathbf{r} - \frac{\mathbf{r}_i(t)+\mathbf{r}_j(t)}{2}\right)\delta\left(2\mathbf{l} - (\mathbf{r}_j(t) - \mathbf{r}_i(t))\right)\right)\Bigg]\\
&+\frac{(v^j_\alpha - v^i_\alpha) - (\frac{l^{ji}_\alpha}{l_{ji}}\frac{l^{ji}_\gamma}{l_{ji}})(v^j_\gamma - v^i_\gamma)}{l_{ji}}n_{\rm p}^{ij}\left(\frac{l^{ji}_\beta}{2}\delta\left(\mathbf{r} - \frac{\mathbf{r}_i(t)+\mathbf{r}_j(t)}{2}\right)\delta\left(2\mathbf{l} - (\mathbf{r}_j(t) - \mathbf{r}_i(t))\right)\right)\\
&+\left(\frac{k_{\rm on}e^{-l_{ji}/l_0}}{2\pi l_0^2}\frac{l^{ji}_\alpha}{l_{ji}} - k_{\rm off}n_{\rm p}^{ij}\frac{l^{ji}_\alpha}{l_{ji}}\right)\left(\frac{l^{ji}_\beta}{2}\delta\left(\mathbf{r} - \frac{\mathbf{r}_i(t)+\mathbf{r}_j(t)}{2}\right)\delta\left(2\mathbf{l} - (\mathbf{r}_j(t) - \mathbf{r}_i(t))\right)\right)\Bigg\rangle\\
= f_{\rm p}\int 4d^2l\Bigg\{\sum_{i\neq j}\Bigg[&\frac{l_\alpha}{l}\left\langle n_{\rm p}^{ij}\delta\left(\mathbf{r} - \frac{\mathbf{r}_i(t)+\mathbf{r}_j(t)}{2}\right)\delta\left(2\mathbf{l} - (\mathbf{r}_j(t) - \mathbf{r}_i(t))\right)\right\rangle\frac{v_\beta(\mathbf{r}+\mathbf{l}) - v_\beta(\mathbf{r}-\mathbf{l})}{2}\Bigg]
\end{aligned}$$

$$-\partial_\gamma\left[\frac{l_\alpha l_\beta}{l}\left\langle n_{\rm p}^{ij}\delta\left(\mathbf{r}-\frac{\mathbf{r}_i(t)+\mathbf{r}_j(t)}{2}\right)\delta\left(2\mathbf{l}-(\mathbf{r}_j(t)-\mathbf{r}_i(t))\right)\right\rangle\frac{v_\gamma(\mathbf{r}+\mathbf{l})+v_\gamma(\mathbf{r}-\mathbf{l})}{2}\right]$$
$$+\left[\frac{l_\beta}{l}\left\langle n_{\rm p}^{ij}\delta\left(\mathbf{r}-\frac{\mathbf{r}_i(t)+\mathbf{r}_j(t)}{2}\right)\delta\left(2\mathbf{l}-(\mathbf{r}_j(t)-\mathbf{r}_i(t))\right)\right\rangle\frac{v_\alpha(\mathbf{r}+\mathbf{l})-v_\alpha(\mathbf{r}-\mathbf{l})}{2}\right]$$
$$-\left[\frac{l_\alpha l_\beta l_\gamma}{l^3}\left\langle n_{\rm p}^{ij}\delta\left(\mathbf{r}-\frac{\mathbf{r}_i(t)+\mathbf{r}_j(t)}{2}\right)\delta\left(2\mathbf{l}-(\mathbf{r}_j(t)-\mathbf{r}_i(t))\right)\right\rangle\frac{v_\gamma(\mathbf{r}+\mathbf{l})-v_\gamma(\mathbf{r}-\mathbf{l})}{2}\right]$$
$$+\left[\frac{k_{\rm on}e^{-2l/l_0}}{2\pi l_0^2}\frac{l_\alpha l_\beta}{l}\left\langle\delta\left(\mathbf{r}-\frac{\mathbf{r}_i(t)+\mathbf{r}_j(t)}{2}\right)\delta\left(2\mathbf{l}-(\mathbf{r}_j(t)-\mathbf{r}_i(t))\right)\right\rangle\right.$$
$$\left.\left.-k_{\rm off}\frac{l_\alpha l_\beta}{l}\left\langle n_{\rm p}^{ij}\delta\left(\mathbf{r}-\frac{\mathbf{r}_i(t)+\mathbf{r}_j(t)}{2}\right)\delta\left(2\mathbf{l}-(\mathbf{r}_j(t)-\mathbf{r}_i(t))\right)\right\rangle\right]\right\}$$
$$\simeq f_{\rm p}\Bigg[N_{\alpha\gamma}\partial_\gamma v_\beta-\partial_\gamma(v_\gamma N_{\beta\alpha})+N_{\beta\gamma}\partial_\gamma v_\alpha-\frac{C}{Nl_0}N_{\alpha\beta}N_{\epsilon\gamma}\partial_\epsilon v_\gamma$$
$$+4k_{\rm on}\int d^2l\left(\frac{l_\beta l_\alpha}{l}\frac{e^{-2l/l_0}}{2\pi l_0^2}n(\mathbf{r}+\mathbf{l})n(\mathbf{r}-\mathbf{l})\right)-k_{\rm off}N_{\alpha\beta}\Bigg].\qquad\text{(S15)}$$

To arrive at this simple expression, we used the coarse-graining rules (eq. (S5) and (S6)), Taylor expansion for small $\mathbf{l}$, and the exchanging $l_\alpha\to l_\alpha^{ji}$ due to the delta function $\delta\left(2\mathbf{l}-(\mathbf{r}_j(t)-\mathbf{r}_i(t))\right)$. It is easy to see that the first line in the final result originates in the relative motion of bacteria, while the second line stems from the pili dynamics. The last term in the first line deserves a bit more attention. The term $\int 4d^2l\sum_{i\neq j}\left\langle\frac{l_\alpha l_\beta l_\gamma l_\epsilon}{l^3}n_{\rm p}^{ij}\delta\left(\mathbf{r}-\frac{\mathbf{r}_i(t)+\mathbf{r}_j(t)}{2}\right)\delta\left(2\mathbf{l}-(\mathbf{r}_j(t)-\mathbf{r}_i(t))\right)\right\rangle$ is the fourth rank tensor. The time evolution of this fourth rank tensor will couple with higher rank tensors. Therefore, an additional closure approximation rule is needed which is a common point in this type of derivation. The additional rules we introduce for this fourth rank nematic tensor are in fact the Doi closure [3]:

$$\int 4d^2l\sum_{i\neq j}\left\langle\frac{l_\alpha l_\beta l_\gamma l_\epsilon}{l^2}n_{\rm p}^{ij}\delta\left(\mathbf{r}-\frac{\mathbf{r}_i(t)+\mathbf{r}_j(t)}{2}\right)\delta\left(2\mathbf{l}-(\mathbf{r}_j(t)-\mathbf{r}_i(t))\right)\right\rangle$$
$$\simeq\int 4d^2l\sum_{i\neq j}\left\langle\frac{l_\alpha^{ji}l_\beta^{ji}}{l_{ji}}n_{\rm p}^{ij}\delta\left(\mathbf{r}-\frac{\mathbf{r}_i(t)+\mathbf{r}_j(t)}{2}\right)\delta\left(2\mathbf{l}-(\mathbf{r}_j(t)-\mathbf{r}_i(t))\right)\right\rangle\cdot$$
$$\int 4d^2l\sum_{i\neq j}\left\langle\frac{l_\gamma^{ji}l_\epsilon^{ji}}{l_{ji}}n_{\rm p}^{ij}\delta\left(\mathbf{r}-\frac{\mathbf{r}_i(t)+\mathbf{r}_j(t)}{2}\right)\delta\left(2\mathbf{l}-(\mathbf{r}_j(t)-\mathbf{r}_i(t))\right)\right\rangle\cdot$$
$$\left(\int 4d^2l\sum_{i\neq j}\left\langle n_{\rm p}^{ij}\delta\left(\mathbf{r}-\frac{\mathbf{r}_i(t)+\mathbf{r}_j(t)}{2}\right)\delta\left(2\mathbf{l}-(\mathbf{r}_j(t)-\mathbf{r}_i(t))\right)\right\rangle\right)^{-1}$$
$$=\frac{C}{N}N_{\beta\alpha}N_{\epsilon\gamma},\qquad\text{(S16)}$$

and the weak approximation:

$$\sum_{i\neq j}\left\langle\frac{A_{ij}}{B_{ij}}\delta\Big(\mathbf{r}-\mathbf{r}_i(t)\Big)\delta\Big(\mathbf{r}'-\mathbf{r}_j(t)\Big)\right\rangle$$

$$\simeq \frac{\sum_{i\neq j}\left\langle A_{ij}\delta\Big(\mathbf{r}-\mathbf{r}_i(t)\Big)\delta\Big(\mathbf{r}'-\mathbf{r}_j(t)\Big)\right\rangle \sum_{i\neq j}\left\langle \delta\Big(\mathbf{r}-\mathbf{r}_i(t)\Big)\delta\Big(\mathbf{r}'-\mathbf{r}_j(t)\Big)\right\rangle}{\sum_{i\neq j}\left\langle B_{ij}\delta\Big(\mathbf{r}-\mathbf{r}_i(t)\Big)\delta\Big(\mathbf{r}'-\mathbf{r}_j(t)\Big)\right\rangle}$$
$$= n(\mathbf{r},t)n(\mathbf{r}',t)\frac{A(\mathbf{r},\mathbf{r}',t)}{B(\mathbf{r},\mathbf{r}',t)}, \tag{S17}$$

where we exchanged $l_\alpha \to l_\alpha^{ji}$ due to the delta function $\delta\left(2\mathbf{l}-(\mathbf{r}_j(t)-\mathbf{r}_i(t))\right)$ and the additional integration stems from the equations (S12) and (S13). The factor $C$ is the phenomenological coefficient, which may serve to improve the approximation, but which we set to 1 in this article. The additional field $N$ is defined as:

$$N = \int d^2r' \sum_{i\neq j}\left\langle n_{\mathrm{p}}^{ij}\delta\Big(\mathbf{r}-\mathbf{r}_i(t)\Big)\delta\Big(\mathbf{r}'-\mathbf{r}_j(t)\Big)\right\rangle$$
$$\simeq \int 4d^2l \sum_{i\neq j}\left\langle n_{\mathrm{p}}^{ij}\delta\left(\mathbf{r}-\frac{\mathbf{r}_i(t)+\mathbf{r}_j(t)}{2}\right)\delta\left(2\mathbf{l}-(\mathbf{r}_j(t)-\mathbf{r}_i(t))\right)\right\rangle, \tag{S18}$$

where the approximation is due to the $\mathbf{l}_{ji} = -\mathbf{l}_{ij}$. The additional field $N$ corresponds to the density of the total number of bound pili at position $\mathbf{r}$.

The combination of these two additional rules gives the fourth rank nematic tensor $\int 4d^2l \sum_{i\neq j}\left\langle \frac{l_\alpha l_\beta l_\gamma l_\epsilon}{l^3} n_{\mathrm{p}}^{ij}\delta\left(\mathbf{r}-\frac{\mathbf{r}_i(t)+\mathbf{r}_j(t)}{2}\right)\delta\left(2\mathbf{l}-(\mathbf{r}_j(t)-\mathbf{r}_i(t))\right)\right\rangle$ as:

$$\int 4d^2l \sum_{i\neq j}\left\langle \frac{l_\alpha l_\beta l_\gamma l_\epsilon}{l^3} n_{\mathrm{p}}^{ij}\delta\left(\mathbf{r}-\frac{\mathbf{r}_i(t)+\mathbf{r}_j(t)}{2}\right)\delta\left(2\mathbf{l}-(\mathbf{r}_j(t)-\mathbf{r}_i(t))\right)\right\rangle$$
$$= \frac{C}{Nl_0}N_{\beta\alpha}N_{\epsilon\gamma}, \tag{S19}$$

The time evolution of the total number density of bound pili $N(\mathbf{r},t)$ can be written by the same procedure as in equation (S15):

$$\partial_t N = \int 4d^2l \sum_{i\neq j}\left\{\left\langle n_{\mathrm{p}}^{ij}\partial_t\left[\delta\left(\mathbf{r}-\frac{\mathbf{r}_i(t)+\mathbf{r}_j(t)}{2}\right)\delta\left(2\mathbf{l}-(\mathbf{r}_j(t)-\mathbf{r}_i(t))\right)\right]\right\rangle + \left\langle \delta\left(\mathbf{r}-\frac{\mathbf{r}_i(t)+\mathbf{r}_j(t)}{2}\right)\delta\left(2\mathbf{l}-(\mathbf{r}_j(t)-\mathbf{r}_i(t))\right)\partial_t n_{\mathrm{p}}^{ij}\right\rangle\right\}$$
$$\simeq -\nabla\cdot(\mathbf{v}N) + 4k_{\mathrm{on}}\int_0^\infty\int_0^{2\pi} l\,dl\,d\theta \frac{e^{-2l/l_0}}{2\pi l_0^2}n(\mathbf{r}+\mathbf{l})n(\mathbf{r}-\mathbf{l}) - k_{\mathrm{off}}N. \tag{S20}$$

#### C. The passive stress tensor

The passive stress tensor can be written as:

$$\underline{\underline{\sigma}}^{\mathrm{p}} = \int d^2r' \sum_{i\neq j}\left\langle -\frac{\partial U_{ij}}{\partial \mathbf{r}_i}\frac{l_{ji}}{2}\hat{\mathbf{l}}_{ji}\delta\Big(\mathbf{r}'-\mathbf{r}_j(t)-\frac{\mathbf{l}_{ji}}{2}\Big)\delta\Big(\mathbf{r}-\mathbf{r}_i(t)-\frac{\mathbf{l}_{ji}}{2}\Big)\right\rangle, \tag{S21}$$

where $U_{ij} = U(\mid \mathbf{r}_i - \mathbf{r}_j \mid)$ is the potential energy between bacteria $i$ and $j$. For the short range potential with the interaction length $\lambda$, the integration is almost zero at $l > \lambda$ and the passive stress tensor becomes:

$$\underline{\underline{\sigma}}^{\mathrm{p}} \simeq \int_{l \lessapprox \lambda} 4d^2 l \sum_{i \neq j} \left\langle -\frac{\partial U_{ij}}{\partial \mathbf{r}_i} \frac{l_{ji}}{2} \hat{\mathbf{l}}_{ji} \delta \left( \mathbf{r} - \frac{\mathbf{r}_i(t) + \mathbf{r}_j(t)}{2} \right) \delta \left( 2\mathbf{l} - (\mathbf{r}_j(t) - \mathbf{r}_i(t)) \right) \right\rangle$$
$$\simeq \int_{l \lessapprox \lambda} 4d^2 l W(\mathbf{r}, \mathbf{l}) \hat{\mathbf{l}}\hat{\mathbf{l}} n(\mathbf{r} + \mathbf{l}) n(\mathbf{r} - \mathbf{l}), \tag{S22}$$

where $W(\mathbf{r}, \mathbf{l})\hat{\mathbf{l}}\hat{\mathbf{l}}$ is the average virial term $-\frac{\partial U_{ij}}{\partial \mathbf{r}_i} \frac{l_{ji}}{2} \hat{\mathbf{l}}_{ji}$ defined by the coarse-grained rules (eq. (S6)):

$$W(\mathbf{r}, \mathbf{l})\hat{\mathbf{l}}\hat{\mathbf{l}} \equiv \frac{\left\langle \sum_{i \neq j} -\frac{\partial U_{ij}}{\partial \mathbf{r}_i} \frac{l_{ji}}{2} \hat{\mathbf{l}}_{ji} \delta \left( \mathbf{r} - \frac{\mathbf{r}_i(t) + \mathbf{r}_j(t)}{2} \right) \delta \left( 2\mathbf{l} - (\mathbf{r}_j(t) - \mathbf{r}_i(t)) \right) \right\rangle}{\left\langle \sum_{i \neq j} \delta \left( \mathbf{r} - \frac{\mathbf{r}_i(t) + \mathbf{r}_j(t)}{2} \right) \delta \left( 2\mathbf{l} - (\mathbf{r}_j(t) - \mathbf{r}_i(t)) \right) \right\rangle}. \tag{S23}$$

Because of the short range interaction, the bacteria pairs can be assumed to be at the nearby position $\mathbf{r}$ and $W(\mathbf{r}, \mathbf{l})$ is:

$$W(\mathbf{r}, \mathbf{l}) = -\frac{1}{(V n(\mathbf{r}))^2} \left\langle \sum_{i \neq j} -\frac{\partial U_{ij}}{\partial \mathbf{r}_i} \frac{l_{ji}}{2} \hat{\mathbf{l}}_{ji} \delta \left( \mathbf{r} - \frac{\mathbf{r}_i(t) + \mathbf{r}_j(t)}{2} \right) \delta \left( 2\mathbf{l} - (\mathbf{r}_j(t) - \mathbf{r}_i(t)) \right) \right\rangle$$
$$\simeq \frac{3 V n(\mathbf{r})}{(V n(\mathbf{r}))^2} \left\langle \frac{\partial U(l)}{\partial l} \frac{l}{2} \hat{\mathbf{l}}\hat{\mathbf{l}} \right\rangle, \tag{S24}$$

where $V = \int_{l \lessapprox \lambda} 4d^2 l$ is the volume of interest, $Vn$ is the total number of bacteria in the small region of interest, and the factor $3Vn$ results from Delaunay triangulation [4]. Thus, the passive stress tensor becomes:

$$\underline{\underline{\sigma}}^{\mathrm{p}} \simeq \int_{l \lessapprox \lambda} 4d^2 l \frac{3}{2 V n(\mathbf{r})} \left\langle \frac{\partial U(l)}{\partial l} l \hat{\mathbf{l}}\hat{\mathbf{l}} \right\rangle n(\mathbf{r} + \mathbf{l}) n(\mathbf{r} - \mathbf{l})$$
$$\simeq \frac{3}{2} n(\mathbf{r}) \left\langle \frac{\partial U(l)}{\partial l} l \hat{\mathbf{l}}\hat{\mathbf{l}} \right\rangle \Bigg|_{l \to \frac{1}{\sqrt{\pi n}}}. \tag{S25}$$

Here the mid-point rule $\int dx f(x) \simeq V f(\langle x \rangle)$ has been utilized, $\langle \cdots \rangle$ denotes the ensemble average, and we used next nearest neighbor distance $l \sim \frac{1}{\sqrt{\pi n}}$ to evaluate the passive stress tensor.

For further calculations we need to specify the concrete functional form of the potential. Here we choose the following potential:

$$U_{ji}(l_{ji}) = -\frac{E}{3} \frac{l_{ji}^2 \pi}{4 v_{\mathrm{c}}} \exp\left( -\frac{l_{ji}}{\lambda} \right) \log\left( 1 - \frac{4 v_{\mathrm{c}}}{l_{ji}^2 \pi} \right), \tag{S26}$$

where $v_{\mathrm{c}}$ is the volume of a bacterium, $E$ is the repulsion energy, and the factor $\exp\left(-\frac{l_{ji}}{\lambda}\right)$ signals the short rangeness of the potential. For short distance $l_{ji}^2 \sim \frac{4 v_{\mathrm{c}}}{\pi}$ the potential has a logarithmic divergence to model the excluded volume repulsion. We should note here that the particular shape of the potential is of secondary significance. Its purpose is to introduce repulsive, short range force to counteract pili-induced compression. Then the passive stress tensor can be calculated:

$$\underline{\underline{\sigma}}^{\mathrm{p}} \simeq -\left( \frac{n}{1 - n v_{\mathrm{c}}} + \frac{\log(1 - n v_{\mathrm{c}})}{v_{\mathrm{c}}} - \frac{\log(1 - n v_{\mathrm{c}})}{\sqrt{\pi n} v_{\mathrm{c}} \lambda} \right) E \exp\left( -\frac{2}{\lambda \sqrt{n \pi}} \right) \left\langle \hat{\mathbf{l}}\hat{\mathbf{l}} \right\rangle. \tag{S27}$$

Since we are interested in the high density regime, $n \to 1/v_{\mathrm{c}}$, the passive stress tensor can be approximated as:

$$\underline{\underline{\sigma}}^{\mathrm{p}} \simeq -\frac{En}{1-nv_{\mathrm{c}}}\left\langle \hat{\mathbf{l}}\hat{\mathbf{l}}\right\rangle . \tag{S28}$$

If furthermore we assume the bacterial distribution to be isotropic at any instance, the traceless part of the stress tensor vanishes and thus leads to the stress tensor that contains the bulk part (diagonal part) only:

$$\begin{aligned}\underline{\underline{\sigma}}^{\mathrm{p}} &= -\frac{En}{1-nv_{\mathrm{c}}}\left\langle \hat{\mathbf{l}}\hat{\mathbf{l}} - \frac{1}{2}\,\mathrm{Tr}[\hat{\mathbf{l}}\hat{\mathbf{l}}]\underline{\underline{I}} + \frac{1}{2}\,\mathrm{Tr}[\hat{\mathbf{l}}\hat{\mathbf{l}}]\underline{\underline{I}}\right\rangle \\ &= -\frac{E' v_{\mathrm{c}} n}{1-nv_{\mathrm{c}}}\underline{\underline{I}}, \end{aligned} \tag{S29}$$

where $E' = \frac{E}{2v_{\mathrm{c}}}$ can be explained as the bulk elastic modulus and $\underline{\underline{I}}$ is the identity matrix. It is worth to mention that the factor $\frac{n}{1-nv_{\mathrm{c}}}$ is in fact the formula for pressure of the van der Waals gas without attraction interaction. This is one of the simplest ways to account for excluded volume interactions in the passive stress tensor, while other functional forms of the excluded volume interactions would only change the physical behavior quantitatively but not qualitatively.

#### D. The stress relaxation

This system with pili-induced interactions exhibits viscoelastic properties, which can be shown by combining the equation (S14) and the time derivative of the active stress tensor (eq. (S15)):

$$\begin{aligned}(1+\tau\frac{D}{Dt})\sigma^{\mathrm{a}}_{\alpha\beta} &= \tau f_{\mathrm{p}} N_{\alpha\gamma}\partial_\gamma v_\beta - \tau f_{\mathrm{p}} N_{\beta\alpha}\partial_\gamma v_\gamma + \tau f_{\mathrm{p}} N_{\beta\gamma}\partial_\gamma v_\alpha \\ &\quad - \frac{f_{\mathrm{p}}\tau}{N l_0} N_{\beta\alpha} N_{\epsilon\gamma}\partial_\epsilon v_\gamma + 4\tau f_{\mathrm{p}} k_{\mathrm{on}} \int d^2 l \frac{l_\beta l_\alpha}{l}\frac{e^{-2l/l_0}}{2\pi l_0^2}\left(n(\mathbf{r}+\mathbf{l})n(\mathbf{r}-\mathbf{l})\right),\end{aligned}$$

where $\tau = k_{\mathrm{off}}^{-1}$ is the relaxation time, and $\frac{D}{Dt}$ is the total time derivative. Since the active stress tensor is a symmetric tensor ($\sigma^{\mathrm{a}}_{\alpha\beta} = \sigma^{\mathrm{a}}_{\beta\alpha}$), the effective viscosity $\eta^{eff}_{\alpha\beta\gamma\epsilon}$ can be written in a 3-by-4 matrix form:

$$\begin{bmatrix} \eta^{eff}_{xxxx} & \eta^{eff}_{xxyy} & \eta^{eff}_{xxxy} & \eta^{eff}_{xxyx} \\ \eta^{eff}_{yyxx} & \eta^{eff}_{yyyy} & \eta^{eff}_{yyxy} & \eta^{eff}_{yyyx} \\ \eta^{eff}_{xyxx} & \eta^{eff}_{xyyy} & \eta^{eff}_{xyxy} & \eta^{eff}_{xyyx} \end{bmatrix} =$$

$$= \tau f_{\mathrm{p}} \begin{bmatrix} N_{xx} - \frac{N_{xx}N_{xx}}{Nl_0} & -N_{xx} - \frac{N_{xx}N_{yy}}{Nl_0} & 2N_{xy} - \frac{N_{xx}N_{xy}}{Nl_0} & -\frac{N_{xx}N_{xy}}{Nl_0} \\ -N_{yy} - \frac{N_{yy}N_{p,xx}}{Nl_0} & N_{yy} - \frac{N_{yy}N_{yy}}{Nl_0} & -\frac{N_{yy}N_{xy}}{Nl_0} & 2N_{xy} - \frac{N_{yy}N_{xy}}{Nl_0} \\ -\frac{N_{xy}N_{xx}}{Nl_0} & -\frac{N_{xy}N_{yy}}{Nl_0} & N_{yy} - \frac{N_{xy}N_{xy}}{Nl_0} & N_{xx} - \frac{N_{xy}N_{xy}}{Nl_0} \end{bmatrix}.$$

#### E. The bound pili formation

The contribution from bound pili formation appears in the nematic tensor and the total number of bound pili (equations (S15) and (S18))) via the non-local term $n(\mathbf{r}+\mathbf{l})n(\mathbf{r}-\mathbf{l})$. In order to close the equations, the Taylor expansion is needed, where we expand up to the second order spatial derivatives. The contribution of the bound pili formation in the nematic tensor and the total number of bound pili become respectively:

$$4k_{\text{on}}\int_0^\infty\int_0^{2\pi} l dl d\theta \frac{l_\beta l_\alpha}{l}\frac{e^{-2l/l_0}}{2\pi l_0^2}\left(n(\mathbf{r}+\mathbf{l})n(\mathbf{r}-\mathbf{l})\right)$$
$$\simeq 4k_{\text{on}}\int_0^\infty\int_0^{2\pi} l dl d\theta \frac{l_\beta l_\alpha}{l}\frac{e^{-2l/l_0}}{2\pi l_0^2}\left(n^2-(\mathbf{l}\cdot\nabla n)^2+n(\mathbf{l}\cdot\nabla)^2 n\right) \tag{S30}$$

and

$$4k_{\text{on}}\int_0^\infty\int_0^{2\pi} l dl d\theta \frac{e^{-2l/l_0}}{2\pi l_0^2}\left(n(\mathbf{r}+\mathbf{l})n(\mathbf{r}-\mathbf{l})\right)$$
$$\simeq 4k_{\text{on}}\int_0^\infty\int_0^{2\pi} l dl d\theta \frac{e^{-2l/l_0}}{2\pi l_0^2}\left(n^2-(\mathbf{l}\cdot\nabla n)^2+n(\mathbf{l}\cdot\nabla)^2 n\right). \tag{S31}$$

Because of the exponential factor in the integration, the above equations can be written as:

$$\begin{aligned}&4k_{\text{on}}\int_0^\infty\int_0^{2\pi} l dl d\theta \frac{l_\beta l_\alpha}{l}\frac{e^{-2l/l_0}}{2\pi l_0^2}\left(n^2-(\mathbf{l}\cdot\nabla n)^2+n(\mathbf{l}\cdot\nabla)^2 n\right)\\ &=\frac{k_{\text{on}}}{2}\Bigg\{\begin{bmatrix} l_0 n^2 & 0\\ 0 & l_0 n^2\end{bmatrix}+\begin{bmatrix} -\frac{3l_0^3}{4}\left(3(\partial_x n)^2+(\partial_y n)^2\right) & 0\\ 0 & -\frac{3l_0^3}{4}\left((\partial_x n)^2+3(\partial_y n)^2\right)\end{bmatrix}\\ &+\begin{bmatrix}\frac{3l_0^3}{4}\left(3n\partial_x^2 n+n\partial_y^2 n\right) & \frac{3l_0^3}{2}\left(-(\partial_x n)(\partial_y n)+n\partial_x\partial_y n\right)\\ \frac{3l_0^3}{2}\left(-(\partial_x n)(\partial_y n)+n\partial_x\partial_y n\right) & \frac{3l_0^3}{4}\left(n\partial_x^2 n+3n\partial_y^2 n\right)\end{bmatrix}\Bigg\},\end{aligned} \tag{S32}$$

and

$$4k_{\text{on}}\int_0^\infty\int_0^{2\pi} l dl d\theta \frac{e^{-2l/l_0}}{2\pi l_0^2}\left(n^2-(\mathbf{l}\cdot\nabla n)^2+n(\mathbf{l}\cdot\nabla)^2 n\right)$$
$$=\frac{k_{\text{on}}}{2\pi}\left[2\pi n^2-\pi\frac{3l_0^2}{2}\left((\partial_x n)^2+(\partial_y n)^2\right)+\pi\frac{3l_0^2}{2}\left(n\partial_x^2 n+n\partial_y^2 n\right)\right]. \tag{S33}$$

#### F. The overall time evolution equations

To sum up, the overall continuous fields fully describing the dynamics of aggregates are: $n(\mathbf{r},t)$, $\mathbf{v}(\mathbf{r},t)$, $N_{\alpha\beta}(\mathbf{r},t)$, and $N(\mathbf{r},t)$. The corresponding governing equations are (eq. (S7), (S8), (S15), (S18), (S32), and (S33)):

$$\begin{aligned}\partial_t n &= -\nabla\cdot\Big(\mathbf{v}n\Big)\\ \xi n\mathbf{v} &= f_{\text{p}}\nabla\cdot\underline{\underline{N}}-\nabla\cdot\left(\frac{E' v_{\text{c}} n}{1-nv_{\text{c}}}\underline{\underline{I}}\right)\\ \partial_t N_{\alpha\beta} &= N_{\alpha\gamma}\partial_\gamma v_\beta-\partial_\gamma(v_\gamma N_{\beta\alpha})+N_{\beta\gamma}\partial_\gamma v_\alpha-\frac{1}{Nl_0}N_{\alpha\beta}N_{\epsilon\gamma}\partial_\epsilon v_\gamma\\ &\quad +N^f_{\alpha\beta}-k_{\text{off}}N_{\alpha\beta}\\ \underline{\underline{N^f}} &= \frac{k_{\text{on}}}{2}\Bigg\{\begin{bmatrix} l_0 n^2 & 0\\ 0 & l_0 n^2\end{bmatrix}+\begin{bmatrix} -\frac{3l_0^3}{4}\left(3(\partial_x n)^2+(\partial_y n)^2\right) & 0\\ 0 & -\frac{3l_0^3}{4}\left((\partial_x n)^2+3(\partial_y n)^2\right)\end{bmatrix}\end{aligned}$$

$$+\begin{bmatrix} \frac{3l_0^3}{4}\left(3n\partial_x^2 n + n\partial_y^2 n\right) & \frac{3l_0^3}{2}\left(-(\partial_x n)(\partial_y n) + n\partial_x\partial_y n\right) \\ \frac{3l_0^3}{2}\left(-(\partial_x n)(\partial_y n) + n\partial_x\partial_y n\right) & \frac{3l_0^3}{4}\left(n\partial_x^2 n + 3n\partial_y^2 n\right) \end{bmatrix} \Bigg\}$$

$$\partial_t N = -\nabla\cdot(\mathbf{v}N) + \frac{k_{\rm on}}{2\pi}\left[2\pi n^2 - \pi\frac{3l_0^2}{2}\left((\partial_x n)^2 + (\partial_y n)^2\right) + \pi\frac{3l_0^2}{2}\left(n\partial_x^2 n + n\partial_y^2 n\right)\right] - k_{\rm off}N, \tag{S34}$$

where the time evolution of the nematic tensor $N_{\alpha\beta}$ follows directly from the equation (S15) for the active stress tensor and $N^f_{\alpha\beta}$ denotes the contribution of the newly formed pili to the nematic tensor.

#### G. The linear stability analysis

The onset of the active phase separation can be calculated through the linear stability analysis of the equations (S34) with the adiabatic approximation. Then, the number density equation is written as:

$$\begin{aligned}\partial_t n =& -\frac{1}{\xi}\nabla\cdot\left(\frac{f_{\rm p}}{k_{\rm off}}\nabla\cdot\underline{\underline{N}}^f - \nabla\cdot\left(\frac{E' v_{\rm c} n}{1-nv_{\rm c}}\underline{\underline{I}}\right)\right)\\ =& -\frac{f_{\rm p}k_{\rm on}l_0}{2k_{\rm off}\xi}\left(\partial_x^2+\partial_y^2\right)n^2 + \frac{f_{\rm p}k_{\rm on}}{2k_{\rm off}\xi}\frac{3l_0^3}{4}\partial_x^2\left(3(\partial_x n)^2 + (\partial_y n)^2\right) + \frac{f_{\rm p}k_{\rm on}}{2k_{\rm off}\xi}\frac{3l_0^3}{4}\partial_y^2\left((\partial_x n)^2 + 3(\partial_y n)^2\right)\\ & -\frac{f_{\rm p}k_{\rm on}}{2k_{\rm off}\xi}\frac{3l_0^3}{4}\partial_x^2\left(3n\partial_x^2 n + n\partial_y^2 n\right) - \frac{f_{\rm p}k_{\rm on}}{2k_{\rm off}\xi}\frac{3l_0^3}{4}\partial_y^2\left(n\partial_x^2 n + 3n\partial_y^2 n\right) - \frac{f_{\rm p}k_{\rm on}}{k_{\rm off}\xi}\frac{3l_0^3}{2}\partial_x\partial_y\left(-(\partial_x n)(\partial_y n) + n\partial_x\partial_y n\right)\\ & +\frac{1}{\xi}\left(\partial_x^2+\partial_y^2\right)\left(\frac{E' v_{\rm c} n}{1-nv_{\rm c}}\right). \end{aligned} \tag{S35}$$

Assume the original number density is homogeneous $n_0$ but with small density fluctuation $\delta n(x,y)$, then time evolution of the spatially Fourier transformed ($\nabla \to i\vec{q}$, where $i$ is the imaginary unit) density fluctuation $\delta n(q_x,q_y)$ is written as:

$$\begin{aligned}\partial_t \delta n \simeq& \frac{n_0 l_0 f_{\rm p}k_{\rm on}}{k_{\rm off}\xi}\left(q_x^2+q_y^2\right)\delta n - \frac{f_{\rm p}k_{\rm on}n_0}{2k_{\rm off}\xi}\frac{3l_0^3}{4}q_x^2\left(3q_x^2\delta n + q_y^2\delta n\right) - \frac{f_{\rm p}k_{\rm on}n_0}{2k_{\rm off}\xi}\frac{3l_0^3}{4}q_y^2\left(q_x^2\delta n + 3q_y^2\delta n\right)\\ & -\frac{f_{\rm p}k_{\rm on}n_0}{k_{\rm off}\xi}\frac{3l_0^3}{2}\partial_x^2\partial_y^2\delta n - \frac{1}{\xi}\frac{E' v_{\rm c}}{(1-n_0 v_{\rm c})^2}\left(q_x^2+q_y^2\right)\delta n.\end{aligned}$$

Since all the parameters $f_{\rm p}$, $k_{\rm on}$, $k_{\rm off}$, $l_0$, $\xi$, and $n_0$ are positive, the large scale (small $|\,\vec{q}\,|$) structure will emerge when $-\frac{1}{\xi}\frac{E' v_{\rm c}}{(1-n_0v_{\rm c})^2} + \frac{n_0 l_0 f_{\rm p} k_{\rm on}}{k_{\rm off}\xi} > 0$. This indicates the onset of the phase separation.

#### H. Dimensional analysis

The dimensional analysis can be done by using Buckingham $\pi$ theorem for each equation in the above system (equation (S34)). For the force balance equation, the independent variables are: $f_{\rm p}$, $l_0$, $E'$, $v_{\rm c} = \pi R^2$, and $\xi$, and there are three distinct dimensions: time, length, and force. We choose the unit of length $R$, time $\frac{\xi R}{f_{\rm p}}$, and force $f_{\rm p}$, and generate the non-dimensional variables of pili length $\frac{l_0}{R}$ and the excluded volume force $\frac{E'\pi R}{f_{\rm p}}$. For the nematic tensor dynamics, the independent variables are: $k_{\rm on}$, $k_{\rm off}$, and $l_0$, and there are two distinct dimensions: time and length. By choosing the unit of length $l_0$, and time $k_{\rm off}^{-1}$, we derive the non-dimensional variable $\frac{k_{\rm on}}{k_{\rm off}l_0^2}$ corresponding to steady state bound pili density. The equation for the total number of bound pili dynamics gives the same non-dimensional variable as the nematic tensor dynamics. By combining dimensional analysis of the force balance equation and the nematic tensor dynamics, we chose the unit of length to be $R$, the unit of time to be $\frac{\xi R}{f_{\mathrm{p}}}$, and the unit of force to be $f_{\mathrm{p}}$, leading to four non-dimensional variables: $\frac{l_0}{R}$, $\frac{E'\pi R}{f_{\mathrm{p}}}$, $\frac{k_{\mathrm{on}}}{k_{\mathrm{off}}R^2}$, and $\frac{k_{\mathrm{off}}\xi R}{f_{\mathrm{p}}}$. The last term quantifies how many pili detachment evernts happen while the bacterium travels its own radius under the motion of pili force $f_{\mathrm{p}}$.

#### I. The conservation of the total number of bound pili

In the main text, we assume the total number of bound pili is unlimited; however, in reality, the total number of bound pili is often bounded [5]. Assume each bacterium can form $B$ bound pili, the microscopic time evolution of the number of bound pili then is:

$$\frac{d}{dt}n_{\mathrm{p}}^{ji} = \frac{k_{\mathrm{on}}e^{-\frac{l_{ji}}{l_0}}}{2\pi l_0^2}\left(B - \sum_k n_p^{ki}\right)\left(B - \sum_k n_p^{kj}\right) - k_{\mathrm{off}}n_{\mathrm{p}}^{ji}, \tag{S36}$$

and the corresponding continuous fields of the nematic tensor $N_{p,\alpha\beta}$ and and total number of bound pili $N$ are:

$$\partial_t N_{\alpha\beta} = N_{\alpha\gamma}\partial_\gamma v_\beta - \partial_\gamma(v_\gamma N_{\beta\alpha}) + N_{\beta\gamma}\partial_\gamma v_\alpha - \frac{C}{Nl_0}N_{\alpha\beta}N_{\epsilon\gamma}\partial_\epsilon v_\gamma$$
$$+4k_{\mathrm{on}}\int d^2l\left(\frac{l_\beta l_\alpha}{l}\frac{e^{-2l/l_0}}{2\pi l_0^2}\left(Bn(\mathbf{r}+\mathbf{l}) - N(\mathbf{r}+\mathbf{l})\right)\left(Bn(\mathbf{r}-\mathbf{l}) - N(\mathbf{r}-\mathbf{l})\right)\right) - k_{\mathrm{off}}N_{\alpha\beta} \tag{S37}$$

$$\partial_t N = -\nabla\cdot(\mathbf{v}N) + 4k_{\mathrm{on}}\int_0^\infty\int_0^{2\pi} ldld\theta\frac{e^{-2l/l_0}}{2\pi l_0^2}\left(Bn(\mathbf{r}+\mathbf{l}) - N(\mathbf{r}+\mathbf{l})\right)\left(Bn(\mathbf{r}-\mathbf{l}) - N(\mathbf{r}-\mathbf{l})\right) - k_{\mathrm{off}}N, \tag{S38}$$

where the derivation of the newly bound pili formation term is:

$$\int d^2r'\sum_{i\neq j}\frac{k_{\mathrm{on}}e^{-\frac{l_{ji}}{l_0}}}{2\pi l_0^2}\left(B - \sum_{k,\text{for } k\neq i} n_p^{ki}\right)\left(B - \sum_{k,\text{for } k\neq j} n_p^{kj}\right)\delta\Big(\mathbf{r}-\mathbf{r}_i\Big)\delta\left(\mathbf{r}'-\mathbf{r}_j\right)$$
$$= \int d^2r'\sum_{i\neq j}\frac{k_{\mathrm{on}}e^{-\frac{l_{ji}}{l_0}}}{2\pi l_0^2}B^2\delta\Big(\mathbf{r}-\mathbf{r}_i\Big)\delta\left(\mathbf{r}'-\mathbf{r}_j\right)$$
$$-\int d^2r'\int d^2r''\sum_{i\neq j}\frac{k_{\mathrm{on}}e^{-\frac{l_{ji}}{l_0}}}{2\pi l_0^2}B\sum_{k,\text{for } k\neq i} n_p^{ki}\delta\Big(\mathbf{r}-\mathbf{r}_i\Big)\delta\left(\mathbf{r}'-\mathbf{r}_j\right)\delta\left(\mathbf{r}''-\mathbf{r}_k\right)$$
$$-\int d^2r'\int d^2r''\sum_{i\neq j}\frac{k_{\mathrm{on}}e^{-\frac{l_{ji}}{l_0}}}{2\pi l_0^2}B\sum_{k,\text{for } k\neq j} n_p^{kj}\delta\Big(\mathbf{r}-\mathbf{r}_i\Big)\delta\left(\mathbf{r}'-\mathbf{r}_j\right)\delta\left(\mathbf{r}''-\mathbf{r}_k\right)$$
$$+\int d^2r'\int d^2r''\int d^2r'''\sum_{i\neq j}\frac{k_{\mathrm{on}}e^{-\frac{l_{ji}}{l_0}}}{2\pi l_0^2}\sum_{h,\text{for } h\neq j} n_p^{hj}\sum_{k,\text{for } k\neq i} n_p^{kj}\delta\Big(\mathbf{r}-\mathbf{r}_i\Big)\delta\left(\mathbf{r}'-\mathbf{r}_j\right)\delta\left(\mathbf{r}''-\mathbf{r}_k\right)\delta\left(\mathbf{r}'''-\mathbf{r}_h\right)$$

$$= 4k_{\mathrm{on}} \int_0^\infty \int_0^{2\pi} l dl d\theta \frac{e^{-2l/l_0}}{2\pi l_0^2} \left(Bn(\mathbf{r}+\mathbf{l}) - N(\mathbf{r}+\mathbf{l})\right) \left(Bn(\mathbf{r}-\mathbf{l}) - N(\mathbf{r}-\mathbf{l})\right), \tag{S39}$$

where the coarse-grained rules (eq. (S6)) and equation (S18) have been utilized. The newly bound pili network formation term:

$$\int d^2r' \sum_{i\neq j} \frac{l_\beta^{ji} l_\alpha^{ji}}{2l_{ji}} \frac{k_{\mathrm{on}} e^{-\frac{l_{ji}}{l_0}}}{2\pi l_0^2} \left(B - \sum_{k,\mathrm{for}\ k\neq i} n_p^{ki}\right) \left(B - \sum_{k,\mathrm{for}\ k\neq j} n_p^{kj}\right) \delta\Big(\mathbf{r}-\mathbf{r}_i\Big) \delta\left(\mathbf{r}'-\mathbf{r}_j\right)$$

$$= 4k_{\mathrm{on}} \int d^2l \left( \frac{l_\beta l_\alpha}{l} \frac{e^{-2l/l_0}}{2\pi l_0^2} \left(Bn(\mathbf{r}+\mathbf{l}) - N(\mathbf{r}+\mathbf{l})\right) \left(Bn(\mathbf{r}-\mathbf{l}) - N(\mathbf{r}-\mathbf{l})\right) \right) \tag{S40}$$

can be derived through the same method.

---


[1] Maier, B. & Wong, G. C. L. How Bacteria Use Type IV Pili Machinery on Surfaces. *Trends in Microbiology* **23**, 775–788 (2015).

[2] Biais, N., Higashi, D. L., Brujić, J., So, M. & Sheetz, M. P. Force-dependent polymorphism in type IV pili reveals hidden epitopes. *Proceedings of the National Academy of Sciences* **107**, 11358–11363 (2010).

[3] Doi, M. Molecular dynamics and rheological properties of concentrated solutions of rodlike polymers in isotropic and liquid crystalline phases. *Journal of Polymer Science: Polymer Physics Edition* **19**, 229–243 (1981).

[4] Press, W. H., Teukolsky, S. A., Vetterling, W. T. & Flannery, B. P. *Numerical Recipes 3rd Edition: The Art of Scientific Computing* (Cambridge University Press, New York, NY, USA, 2007), 3 edn.

[5] Pönisch, W., Weber, C. A., Juckeland, G., Biais, N. & Zaburdaev, V. Multiscale modeling of bacterial colonies: How pili mediate the dynamics of single cells and cellular aggregates. *New J. Phys.* **19**, 015003 (2017).
